# Defective Nucleotide Catabolism Defines a Subset of Cancers Sensitive to Purine Nucleoside Phosphorylase Inhibition

**DOI:** 10.1101/810093

**Authors:** Evan R. Abt, Vincent Lok, Thuc M. Le, Soumya Poddar, Woosuk Kim, Joseph R. Capri, Gabriel Abril-Rodriguez, Johannes Czernin, Timothy R. Donahue, Thomas Mehrling, Antoni Ribas, Caius G. Radu

## Abstract

Small molecule inhibitors of purine nucleoside phosphorylase (PNP) have been explored as a treatment strategy for leukemia and lymphoma, however, the determinants of response to this class of drugs are incompletely understood. PNP inhibitors impair cell proliferation by preventing catabolism of the nucleoside deoxyguanosine (dG) which induces toxic imbalances amongst intracellular deoxyribonucleotide triphosphate (dNTP) pools following its phosphorylation and trapping by nucleoside kinases. We hypothesized that differential nucleoside uptake or catabolism defines cancer cell lines as either sensitive or resistant to PNP inhibition. Among cancer cell lines we found that T-cell acute lymphoblastic leukemia (T-ALL) cells are uniquely and acutely sensitive to PNP inhibition whereas the B-cell leukemia and solid tumor models are completely resistant. We determined that although the nucleoside scavenging kinase deoxycytidine kinase (dCK) was active in all cells tested, PNP inhibitors only induced dGTP pool increases in sensitive models. By evaluating the expression of key genes involved in nucleotide scavenging, biosynthesis, and phosphohydrolysis in a panel of sensitive and resistant cell lines we found that the dNTP phosphohydrolase SAM histidine aspartate containing protein 1 (SAMHD1) was exclusively expressed in resistant models. Using CRISPR/Cas9 SAMHD1 knockout cell lines, we verified that PNP inhibitor sensitivity is a function of SAMHD1 expression and determined that the pharmacological inhibition of dCK or genetic restoration of SAMHD1 conferred resistance to PNP inhibition. Importantly, we determined that low expression of SAMHD1 is not limited to T-ALL as subset of established and primary solid tumors models are SAMHD1-deficient. These solid tumor models were consistently acutely sensitive to PNP inhibitors which indicates that the utility of PNP inhibitors extends beyond hematological malignancies. Additionally, we found that deoxycytidine (dC) can limit the anti-proliferative effects of PNP inhibitors by competing with dG for phosphorylation by dCK but this effect can be overcome by expression of dC-catabolizing gene cytidine deaminase (CDA). Collectively, these results indicate that SAMHD1, dCK and CDA are critical biomarkers that must be used to stratify patients in clinical trials evaluating pharmacological PNP inhibition.

## INTRODUCTION

A sufficient supply of deoxyribonucleotide triphosphates (dNTPs) is essential for cell growth, survival and proliferation^1, 2^. Thus, the metabolic network that is responsible for the synthesis of dNTPs is tightly regulated by transcriptional, post-translational and allosteric control mechanisms^3, 4^. Cancer cells are particularly reliant on dNTP biosynthesis and this non-oncogene addiction has been leveraged clinically by the use of small molecule inhibitors of rate-limiting metabolic enzymes including dihydroorotate dehydrogenase (DHODH), ribonucleotide reductase (RNR), dihydrofolate reductase (DHFR) and thymidylate synthase (TYMS)^5^. These drugs restrict the ability of cancer cells to synthesize dNTPs and trigger cell cycle arrest: effectively preventing cell proliferation.

Beyond insufficiency, imbalance among dNTP pools results in mutagenesis, altered cell cycle progression or death in specific cell types and can result in developmental defects. Examples include development of mitochondrial neurogastrointestinal encephalopathy syndrome (MNGIE) as a consequence of down-regulation of the thymidine catabolizing gene thymidine phosphorylase (TYMP) and impaired T-cell development in deoxycytidine kinase (dCK) knockout mice resulting from unchecked expansion of dTTP pools in developing hematopoietic cells^6, 7^. Intriguingly, this cell type specific lethality resulting from dNTP imbalance has also been linked to the pathology of severe-combined immuno-deficiency (SCID) a hereditary genetic disorder in humans that is characterized by compromised immune function and premature death. Giblett and colleagues were among the first to identify that inactivating, inherited mutations in the nucleotide catabolic enzymes adenosine deaminase (ADA) and purine nucleoside phosphorylase (PNP) drive the development of SCID^8, 9^. ADA and PNP function sequentially in the catabolism of purine deoxyribonucleosides dA and dG and counter their intracellular accumulation mediated by deoxycytidine kinase dCK^4^. For reasons not well understood defective dG catabolism results in remarkably selective T-cell defects, and otherwise normal cellular development, in individuals with PNP-linked SCID^10^. This tissue specificity is thought to be related to the expression patterns of the nucleoside kinases dCK and differential dG uptake^11^.

Decades later, the observations made by Giblett and colleagues were leveraged for the treatment of cancer with the rationale that pharmacological inhibition of ADA or PNP could be an effective strategy for patients affected by T- and B- cell leukemias and lymphomas^12^. In the 1990s, Schramm and colleagues spearheaded the effort to develop small molecule PNP inhibitors using their knowledge of the transition state substrate-enzyme structure to rationally design drugs with exceptionally high potency and selectivity^13, 14^. PNP inhibitors prevent the catabolism of deoxyguanosine (dG) to guanine and deoxyribose-1-phosphate thereby allowing the intracellular conversion of dG to dGTP which can prevent *de novo* biosynthesis of pyrimidine dNTPs via allosteric inhibition of RNR^15, 16^. Pharmacological PNP inhibition was found to selectively eradicate T-cell leukemias *in vitro* thus mimicking the SCID phenotype^14^. With these encouraging results, the PNP inhibitor Forodesine (also known as BCX-1777 or Immucillin H) progressed to clinical trials for the treatment of relapsed/refectory T- and B-cell leukemias and lymphomas and received approval for the treatment of peripheral T-cell lymphoma in Japan in 2017^17, 18^. Despite excellent tolerability and pharmacodynamic properties in humans (evidenced by plasma accumulation of the PNP substrate dG and depletion in the product of dG catabolism, uric acid) efficacy was observed only in a subset of patients.

It remains an open challenge to identify biomarkers predictive of PNP inhibitor response in cancer. Multiple markers such as ATM, p53, ZAP-70, HGPRT and various nucleoside transporters have been studied in this context but alone do not predict sensitivity^19, 20^. With the goal of identifying metabolic determinants of PNP inhibitor response, we focused our investigation on heterogeneity in nucleoside uptake, accumulation and catabolism across panels of cancer cell lines. We found that the dNTP phosphohydrolase SAM histidine aspartate containing protein 1 (SAMHD1), which catabolizes dGTP into dG and triphosphate, was expressed exclusively in resistant cell lines. We confirmed a role for SAMHD1 in the activity of PNP inhibitors using both loss of function and gain of function genetic models. Additionally, we identified several solid tumor cell lines, including lung cancer and melanoma, which are SAMHD1-deficient and confirmed that these models were sensitive to PNP inhibition. Collectively these results demonstrate that SAMHD1 activity and not disease type *per se* defines PNP inhibitor sensitivity.

## RESULTS

### SAMHD1 mediates response to PNP inhibition in leukemia cell lines

To identify cell line models sensitive and resistant to PNP inhibition, we evaluated the anti-proliferative effects of dG, Forodesine (PNPi), and the combination across a panel of T- and B-acute lymphoblastic leukemia (ALL) cell lines using Cell Titer Glo. Importantly, PNP deficiency alone does not impair cell proliferation or survival *in vitro* and endogenously generated concentrations of its substrate dG must be supplemented to recapitulate the PNP-loss of function phenotype observed *in vivo*. We found that the T-ALL cell lines CCRF-CEM, JURKAT and MOLT4 were eradicated by the combination whereas the B-ALL models NALM6 and RS411 were unaffected (**Figure 1A**). We confirmed these findings using flow cytometry cell cycle analysis in which we observed an increase in the percentage of sub-G1 cells, a maker for cell death, following PNP inhibition only in T-ALL models (**Figure 1B**). Using immunoblot analysis, we found that PNP inhibitor treatment only induced DNA damage (evidenced by H2AX_S139_ phosphorylation) and apoptosis (evidenced by cleaved caspase 3 accumulation) in CCRF-CEM and not in NALM6 cells (**Figure 1C**).

**Figure 1.**
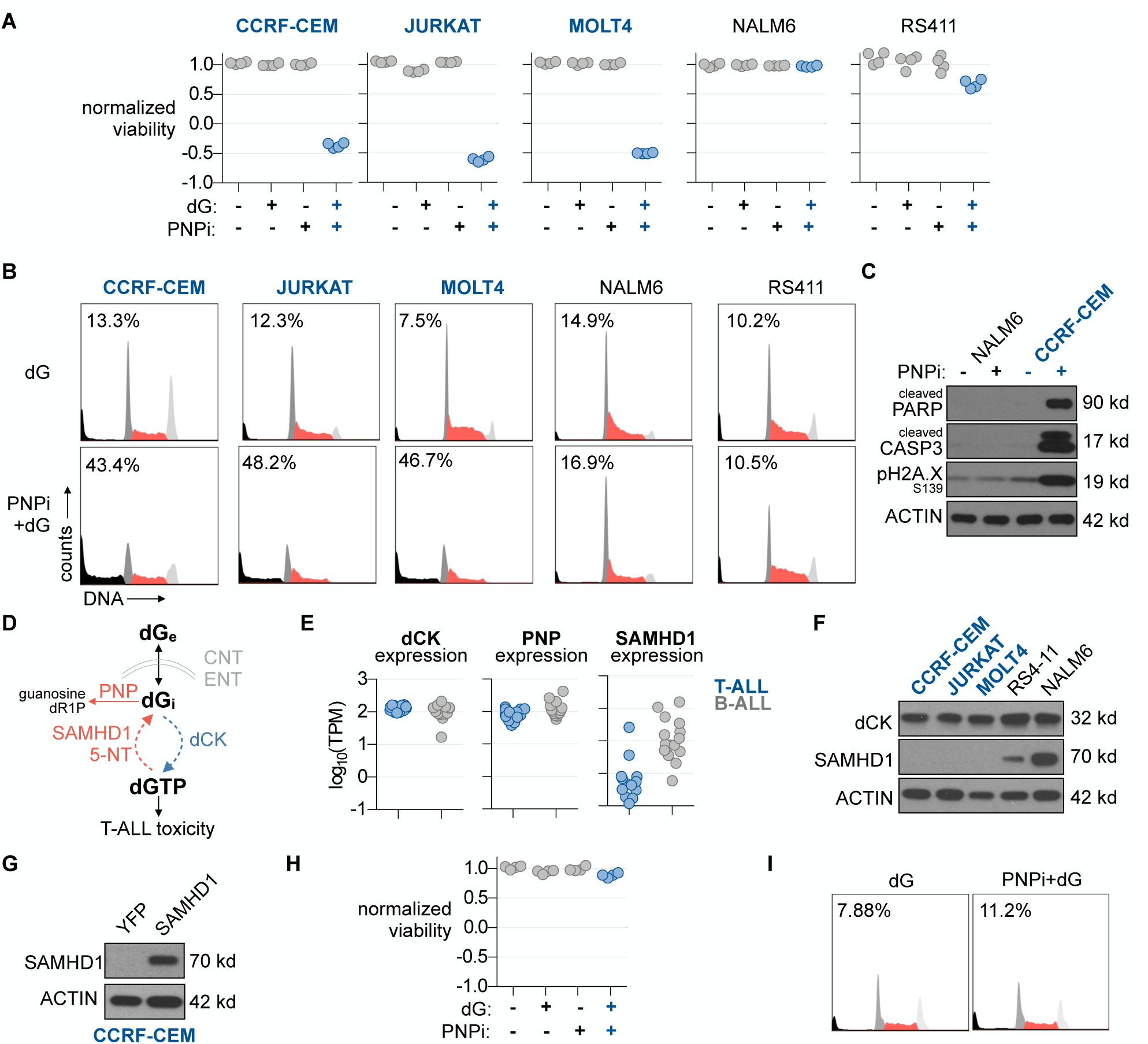
Purine nucleoside phosphorylase (PNP) inhibition is selectively lethal in a subset of acute lymphoblastic leukemia cell lines. (**A**) Cell Titer Glo analysis of acute lymphoblastic leukemia cell line panel treated with 5 µM dG ±1 µM BCX-1777 for 72 h (PNPi, n=4). (**B**) Propidium iodide (PI) cell cycle analysis of acute lymphoblastic leukemia cell lines treated with 5 µM dG ± 1 µM BCX-1777 for 24 h. Insert indicate percentage of Sub-G1 population. (**C**) Immunoblot analysis of NALM6 or CCRF-CEM cells treated with 5 µM dG ± 1 µM BCX-1777 for 24 h. (**D**) Model summarizing metabolic defects potentially explaining sensitivity to PNP inhibitors. (**E**) Expression of dCK, PNP and SAMHD1 across T-ALL cell lines in the Cancer Cell Line Encyclopedia (CCLE) RNAseq dataset. (**F**) Immunoblot analysis of dCK and SAMHD1 expression in T- and B-ALL cell lines. (**G**) Immunoblot validation of SAMHD1 overexpression in CCRF-CEM T-ALL cells. (**H**) Cell Titer Glo analysis of CCRF-CEM YFP control and SAMHD1-over-expressing isogenic cells treated with 5 µM dG ± 1 µM BCX-1777 for 72 h (PNPi, n=4). (**I**) PI cell cycle analysis of CCRF-CEM YFP control and SAMHD1-over-expressing isogenic cells treated with 5 µM dG ± 1 µM BCX-1777 for 24 h. Insert indicate percentage of Sub-G1 population.

We reasoned that heterogeneity in the sensitivity to PNP inhibition could arise from the differential expression of key metabolic genes essential for the synthesis and accumulation of dGTP. These genes include PNP itself, dCK, essential and non-redundant for its ability to phosphorylate dG to dGMP, and SAMHD1, a dNTP phosphohydrolase that catabolizes dGTP to dG (**Figure 1D**). By probing the Cancer Cell Line Encyclopedia (CCLE) gene expression database, we found that dCK and PNP were expressed at similar levels across T- and B-ALL cell line models^21^. In contrast, SAMHD1 was found to be expressed at high levels exclusively in B-ALL cell lines and undetectable in a subset of T-ALL cell lines (**Figure 1E**). Using immunoblot analysis, we confirmed at the protein level that dCK is expressed at similar levels across T- and B-ALL models but SAMHD1 is only detected in B-ALL cell lines (**Figure 1F**). To functionally validate the role of SAMHD1 in the response to PNP inhibition we engineered a CCRF-CEM genetic variant that constitutively expresses human SAMHD1 (**Figure 1G**). We found that expression of SAMHD1 rendered CCRF-CEM cells completely resistant to the anti-proliferative (**Figure 1H**) and cytotoxic effects of PNP (**Figure 1I**).

### SAMHD1-deficient solid tumor models are sensitive to PNP inhibitors

To evaluate the spectrum of SAMHD1 expression across cancers and to determine if low SAMHD1 expression was unique to T-ALL, we extended our analysis to the complete CCLE gene expression dataset (**Figure 2A**). Beyond els, we identified several additional cell lines exhibiting low expression of SAMHD1, including melanoma, lung adenocarcinoma and acute myeloid leukemia cell lines. We hypothesized that low SAMHD1 expression, and not disease type *per se*, predicts response to PNP inhibitors across cancer cell line models. To test this we evaluated the activity of PNPi in the EGFR-mutant, SAMHD1-deficient lung adenocarcinoma cell line HCC827 and found that the combination of dG and PNPi induced cell cycle arrest in this model(**Figure 2B**). We determined that this associated with proliferation inhibition by performing crystal violet analysis of HCC827 cell cultured treated for 7 d (**Figure 2C**).

**Figure 2.**
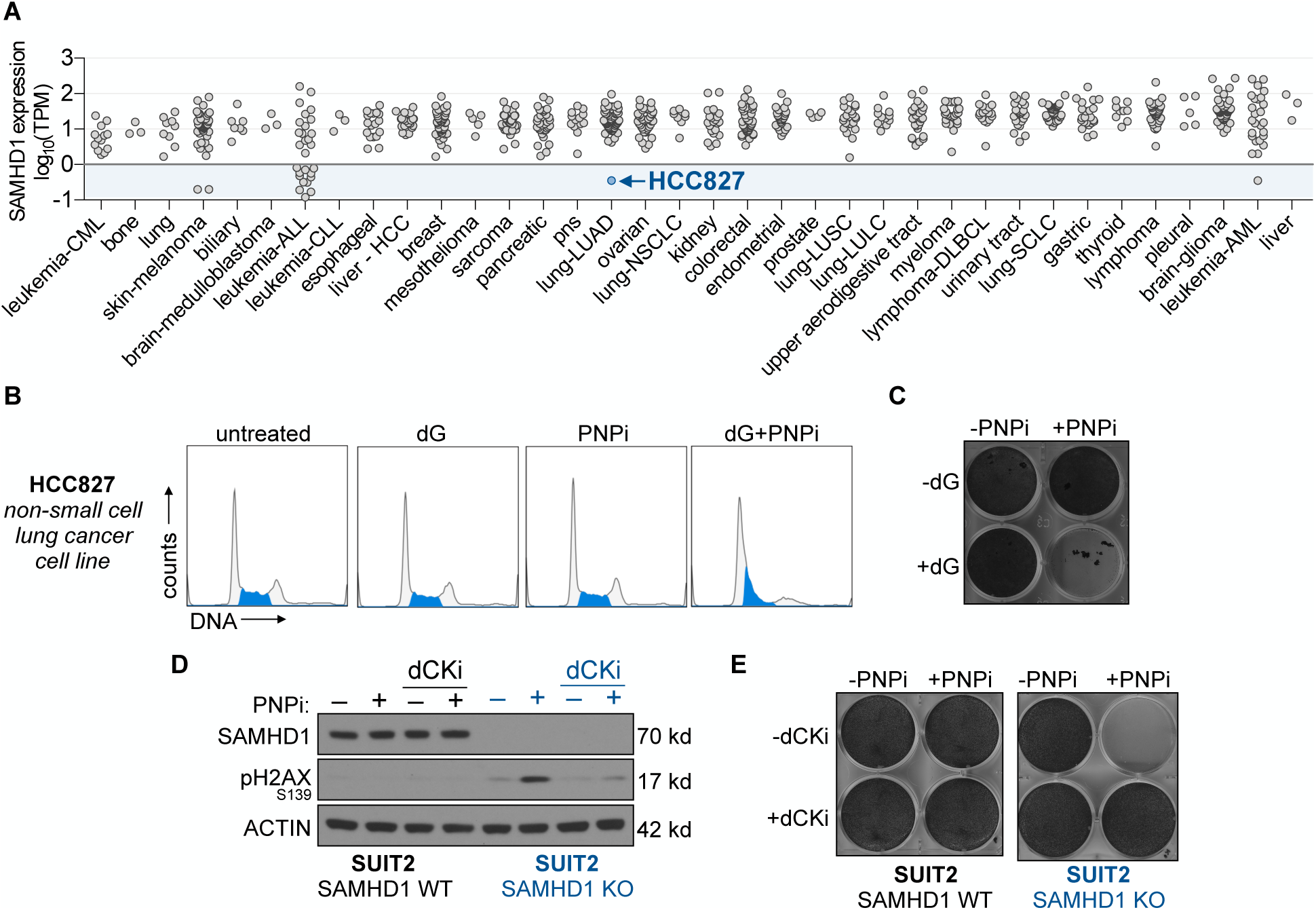
SAMHD1 protects solid tumor-derived cell lines from PNP inhibition. (**A**) SAMHD1 expression in the cancer cell line encyclopedia (CCLE) RNAseq dataset. (**B**) PI cell cycle analysis of the HCC827 non-small cell lung cancer cell line treated with 5 µM dG ± 1 µM BCX-1777 (PNPi) for 24 h. (**C**) Crystal violet analysis of HCC827 cells treated for 7 d with 5 µM dG ± 1 µM PNPi for 7 d. Media was refreshed every 72 h. (**D**) Immunoblot analysis of SUIT2 wild-type (WT) and SAMHD1 knockout (KO) cells treated for 24 h + 5 µM dG ± 1 µM forodesine ± 1 µM (R)-DI-82 (dCKi). (**E**) Crystal violet analysis of SUIT2 WT and SAMHD1 KO cells treated for 7 d with 5 µM dG ± 1 µM PNPi ± 1 µM dCKi for 7 d. Media was refreshed every 72 h.

To functionally validate the role of SAMHD1 in the response of solid tumor cell line models to PNP inhibition, knocked out SAMHD1 in the pancreatic cancer cell line SUIT2 using CRISPR/Cas9. We found that while parental cells were completely resistant to PNPi, SAMHD1 knockout cells exhibited DNA damage in response to PNPi. While dCK is the only cytosolic nucleoside kinase capable of phosphorylating dG, dG is also a substrate for the mitochondria-localized deoxyguanosine kinase (dGK) and SAMHD1 has been shown to generate substrates for dGK in the mitochondria^22^. Furthermore, dCK inactivation has also been associated with resistance to dG-mediated toxicity in PNP-deficient T-lymphoma cells^15^. We found that the lethality of PNP inhibitors be completely prevented by supplementing with a small molecule dCK inhibitor indicating that phosphorylation of dG by dGK does not contribute to the observed lethality at these timepoints (**Figure 2D**)^4^. Additionally, using crystal violet analysis we found that the proliferation of only SAMHD1 knockout cells could be prevented by PNP inhibitors and that this anti-proliferative effect could be completely blocked by a dCK inhibitor (**Figure 2E**). Collectively, these results demonstrate that SAMHD1 expression predicts sensitivity to PNP inhibitors across leukemia and solid tumor cell line models.

### A subset of primary melanoma cell line models are deficient in SAMHD1 expression and are sensitive to PNP inhibitors

To investigate the extent of SAMHD1-deficiency across solid tumor models we profiled the expression of SAMHD1 in a panel of 51 primary melanoma cell lines derived at UCLA (**Figure 3A**). We found that among these models, 2 exhibited undetectable levels of the SAMHD1 transcript (M230, M418) and we confirmed these findings using immunoblot analysis (**Figure 3B**). SAMHD1 function is tightly regulated by transcriptional, post-transcriptional and allosteric control mechanisms and SAMHD1 has been identified as an interferon (IFN)-stimulated gene ^23, 24^. To determine if SAMHD1 expression can be induced in M230 and M418 cells by IFN, we exposed a panel of melanoma cell lines to either type I (IFNβ) or type II (IFNγ) IFN for 24 h and measured SAMHD1 expression using immunoblot analysis (**Figure 3B**). We found that all models tested responded to both type I and type II interferon by increasing the expression of a canonical interferon stimulated gene STAT1, however, SAMHD1 expression was only induced by IFN in cell line models in which SAMHD1 is detectable at baseline. Impaired SAMHD1 expression in M418 and M230 cells could result from mutational inactivation, which has been described in the contested of chronic lymphocytic leukemia, or aberrant promoter methylation which as been described in the context of lung cancer^25, 26^. We confirmed that PNP inhibition selectively inhibited proliferation in SAMHD1-deficient primary melanoma models using crystal violet analysis (**Figure 3C**). Similarly, we observed that PNPi only induced cell cycle arrest in SAMHD1-deficient models (**Figure 3D**).

**Figure 3.**
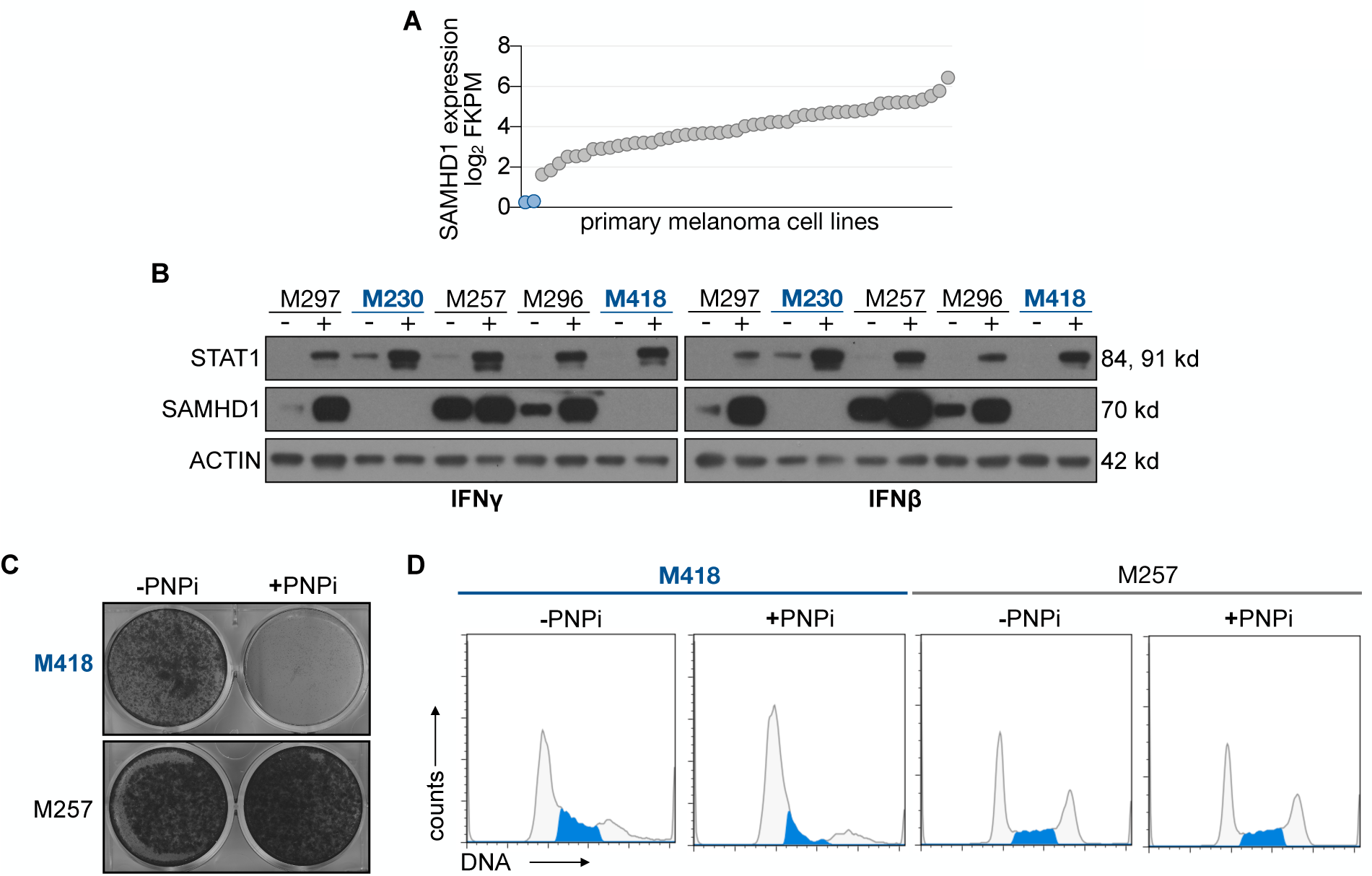
SAMHD1 expression is heterogeneous in primary melanoma cell lines and SAMHD1 deficient cells are sensitive to PNP inhibition. **(A)** RNAseq analysis of SAMHD1 transcript levels in a panel of 51 primary melanoma cell lines. (**B**) Immunoblot analysis of representative melanoma cell lines following 24 h treatment with 100 U/mL IFNβ or 1 ng/mL IFNγ. (**C**) Crystal violet staining following 7 d treatment with 1 µM PNPi (BCX-1777) in the presence of 10 µM dG. (**D**) PI cell cycle analysis following 24 h treatment with 1 µM PNPi in the presence of 10 µM dG.

### dC mitigates the anti-proliferative effects of PNP inhibitors

An additional factor influencing the cytotoxicity of PNP inhibitors is the competition between dG and other deoxyribonucleosides, for dCK (**Figure 4B**). dCK can accept dC, dG and dA as substrates but exhibits an 15-fold higher affinity for dC over the other purine deoxyribonucleosides (BRENDA:EC2.7.1.74). Furthermore, phosphorylation of dCK by ATR further increases its ability to phosphorylate dC while not impacting dA, or dG phosphorylation^27, 28^. In this model, the dC catabolizing enzyme cytidine deaminase (CDA) can promote the activity of PNP inhibitors by eliminating dC and decreasing competition with dG for dCK. To test this model, we evaluated the ability of dC to prevent the anti-proliferative effect of PNP Inhibition in CCEF-CEM YFP control cells and CDA over-expressing cells (**Figure 4B**). In CCRF-CEM YFP cells, PNP inhibition decreased proliferation to 1% of control and supplementation 1 µM dC was sufficient to counteract the effects of PNPi and restored proliferation to 60% of control. Overexpression of CDA prevented this rescue. We have previously reported that plasma dC various greatly across species with levels ranging from 10 nM in human plasma and non-human primates to > 1 µM in rodents (**Figure 4C**; figure adapted from Kim et al. PNAS. 2016). It is conceivable that plasma deoxycytidine can prevent the activity of PNPi and can confound the study of PNP inhibitors in mice. Thus, high-CDA expression should be considered a requisite biomarker for PNP inhibitor sensitivity alongside low-SAMHD1 expression.

**Figure 4.**
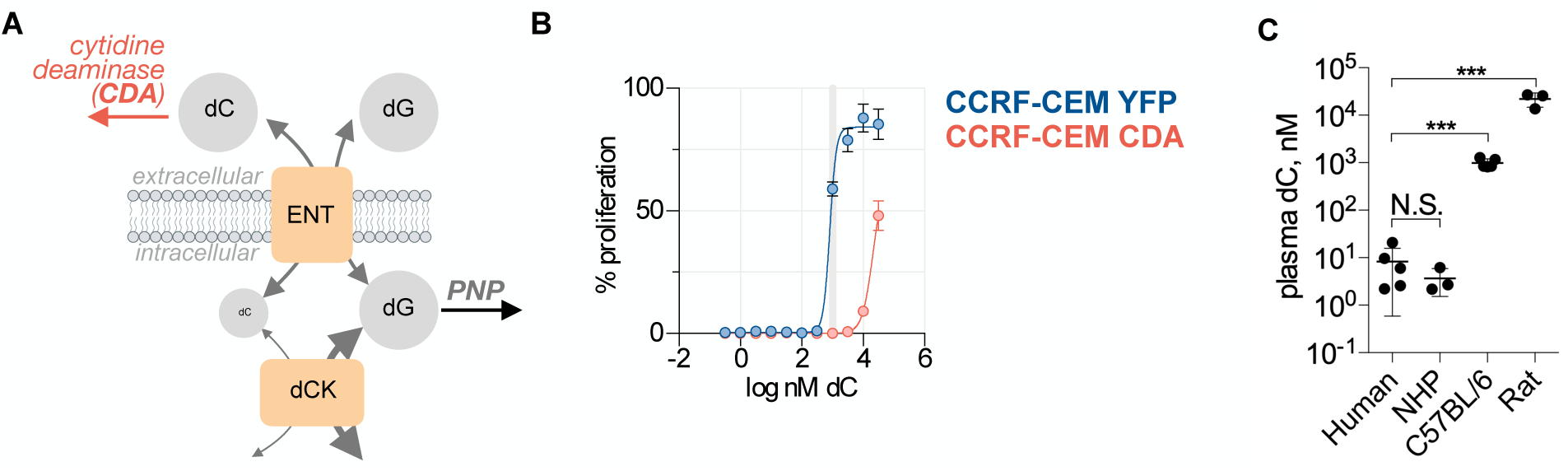
Deoxycytidine competes with dG for phosphorylation by dCK. (**A**) Schematic overview of the competition between dC and dG for dCK. Cytidine deaminase (CDA) breaks down dC and prevents competition with dG. ENT: equilibrate nucleoside transporter. (**B**) Cell Titer Glo analysis of CCRF-CEM YFP and CCRF-CEM CDA over expressing cell lines treated +10 µM dG +1 µM BCX-1777 ± a titration of dC for 72 h.(**C**) Plasma dC in human, non-human primate (NHP), C57BL/6 and rats.

### SAMHD1 prevents the anti proliferative effects of thymidine

Having observed that SAMHD1 can protect cancer cells from dGTP pool imbalance resulting from PNP inhibition we investigated whether SAMHD1 can prevent the anti-proliferative effects of imbalances in other dNTP pools. Elevations in dTTP, dATP or dGTP, but not dCTP, have been linked to impaired cell proliferation which results from allosteric inhibition of RNR^4^. Supplementation of cell cultures with high levels of the pyrimidine nucleoside thymidine is a well established approach to inhibit cell proliferation and synchronize cells in S-phase^29^. Thymidine induced cell cycle arrest is the result of unchecked dTTP pool expansion and allosteric inhibition of CDP reduction by RNR. This “thymidine block” can be completely reversed by supplementation with dC in a dCK-dependent manner. This phenotype extends *in vivo* and is the cause of impaired T-cell development observed in dCK knockout mice^30^. We found that SAMHD1 KO cells exhibited a 200-fold lower thymidine IC_50_ than SAMHD1 WT controls (**Figure 5A**). This selectivity was unique to thymidine and did not extend to other RNR inhibitors, including 3-AP, which function by preventing RNR tyrosyl radical regeneration^4^. Similar selectivity was observed using crystal violet proliferation analysis (**Figure 5B**). Additionally, we found that the induction of pCHEK1_S345_ and pH2A.X_S139_ by dT treatment was enhanced in the SAMHD1 KO cells. Collectively these results indicate that SAMHD1 protects cancer cells from toxic imbalances in both purine and pyrimidine dNTP pools.

**Figure 5.**
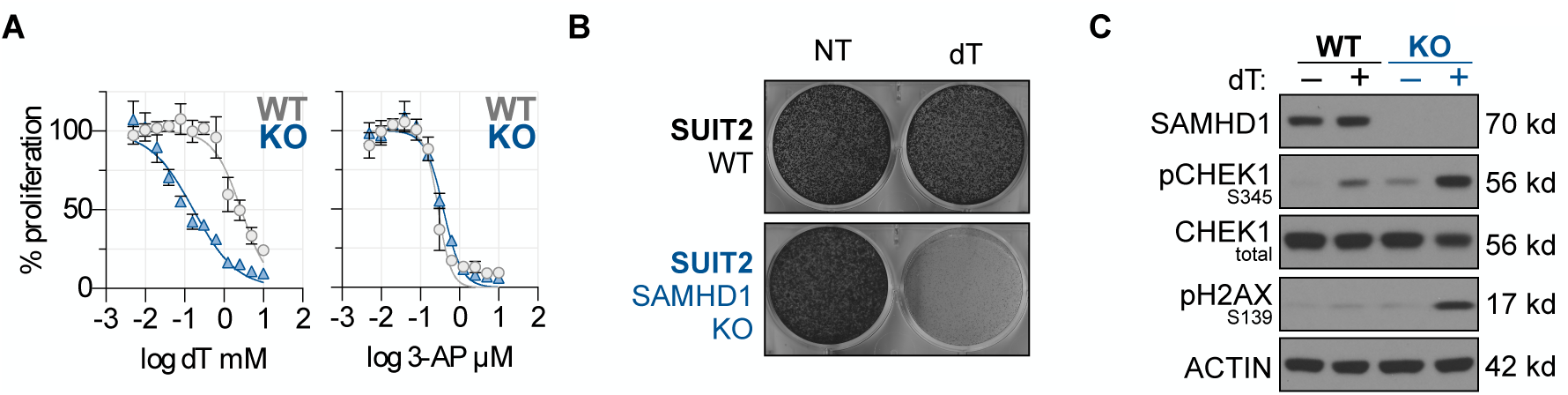
SAMHD1 prevents the toxicity of thymidine. (**A**) Cell Titer Glo analysis of SUIT2 WT and SAMHD1 KO cells treated with a titration of dT or 3-AP for 72 h (mean±SD; n=4). (**B**) Crystal violet analysis of SUIT2 WT and SAMHD1 KO cells treated for 7 d ±100 µM dT. Media was refreshed every 72 h. (**C**) Immunoblot analysis of SUIT2 WT and SAMHD1 KO cells treated for 24 h ±100 µM dT.

### Ulodesine and Forodesine exhibit comparable potency and selectivity

In addition to forodesine an additional small molecule inhibitor of PNP, Ulodesine (also known as BCX-4208) has been developed and evaluated in humans for the treatment of gout^31^. To determine if ulodesine exhibits similar potency and selectivity towards SAMHD1-deficient cancer cells we evaluated inhibition of cell proliferation using Cell Titer Glo (**Figure 6**). We found that both forodesine and ulodesine impaired HCC827 proliferation selectively in the presence of dG with IC50 values of 71 and 21 nM respectively. We expanded this analysis to a panel of cancer cells and found that ulodesine selectively impaired the proliferation of SAMHD1 deficient cancer cell lines JURKAT, CCRF-CEM, and HCC827 while exhibiting negligible activity against SAMHD1-proficient SUIT2 and NALM6 cells (**Figure 7A**). We confirmed a role for SAMHD1 by demonstrating that overexpression of SAMHD1 in CCRF-CEM resulted in resistance and CRISPR/Cas9 knockout of SAMHD1 in SUIT2 cells rendered them sensitive to ulodesine (**Figure 7B**).

**Figure 6.**
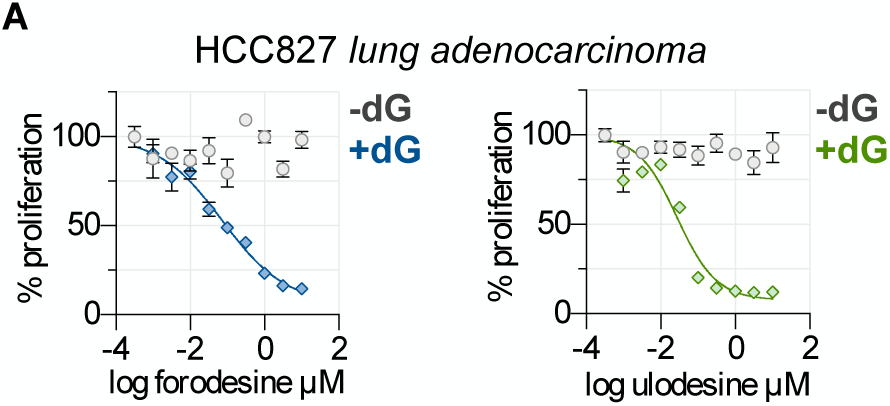
Ulodesine exhibits potency and selectivity similar to forodesine. (**A**) Cell Titer Glo analysis of HCC827 cells treated with a titration of forodesine (BCX-1777) or ulodesine for 72 h ±10 µM dG (mean±SD; n=4).

**Figure 7.**
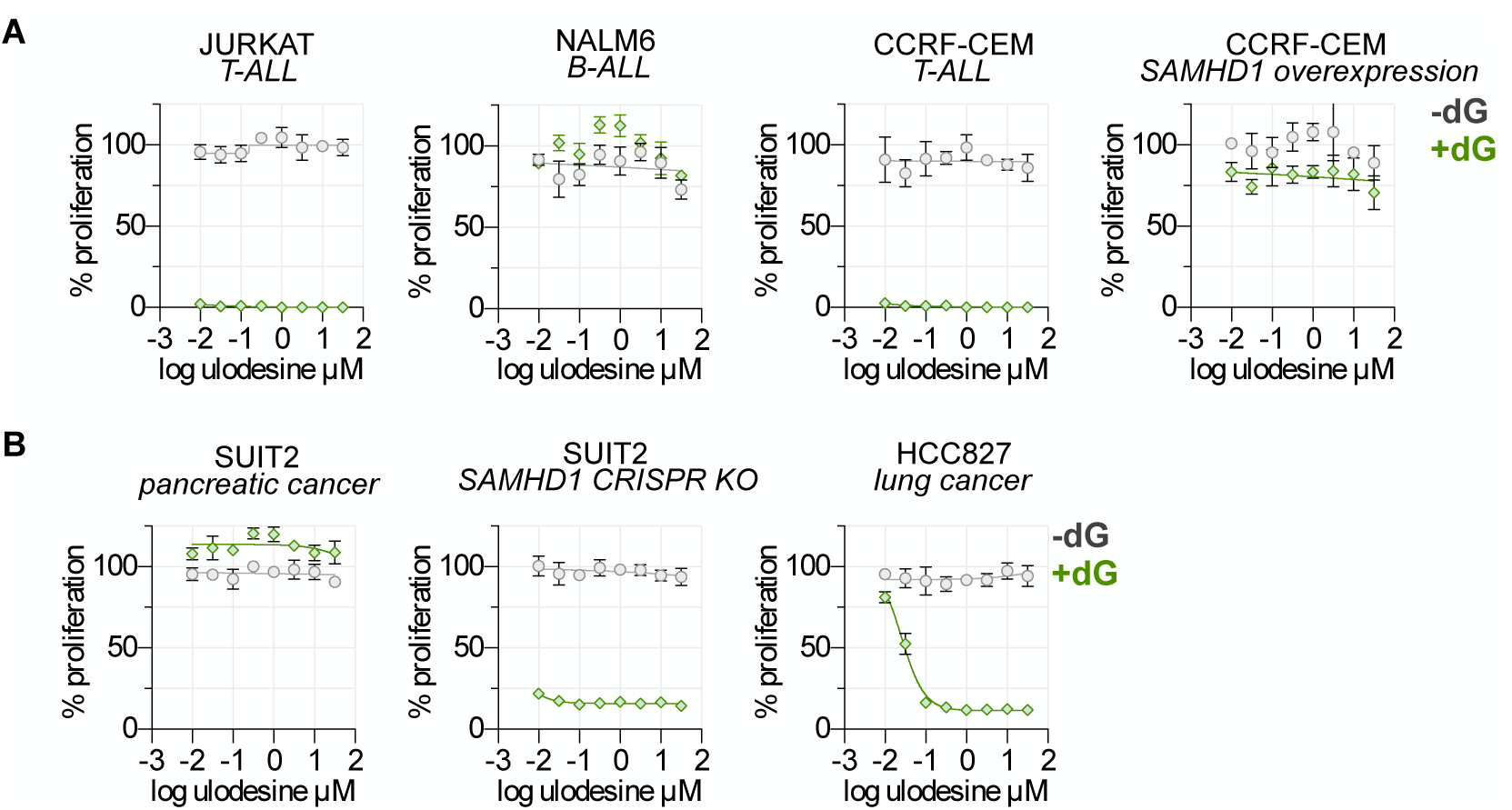
Ulodesine exhibits selective activity in SAMHD1-deficient cancer cell lines. (**A**) Cell Titer Glo analysis of hematopoietic cancer cell lines JURKAT, NALM6, CCRF-CEM and CCRF-CEM SAMHD1 over-expressing cells treated with a titration of ulodesine for 72 h ±1 µM dG (mean±SD; n=4). (**B**) Cell Titer Glo analysis of solid cancer cell lines SUIT2, HCC827 and SUIT2 SAMHD1 CRISPR/Cas9 KO over-expressing cells treated with a titration of ulodesine for 72 h ±5 µM dG (mean±SD; n=4).

### CDA overexpression mitigates competition between dC and dG in HCC827 cells

To determine if CDA can prevent the competition between dC and dG in HCC827 cells we generated CDA over-expressing HCC827 cells using retroviral transduction. We found that HCC827 cells express CDA endogenously which was enhanced with overexpression (**Figure 8A**). We found that dC could prevent the lethality of the dG/ulodesine combination with an EC _50_ of 1.9 µM in control cells. The dC EC _50_ was increased over 11-fold to 21 µM in the CDA over-expressing isogenic cells (**Figures 8A, 8B**).

**Figure 8.**
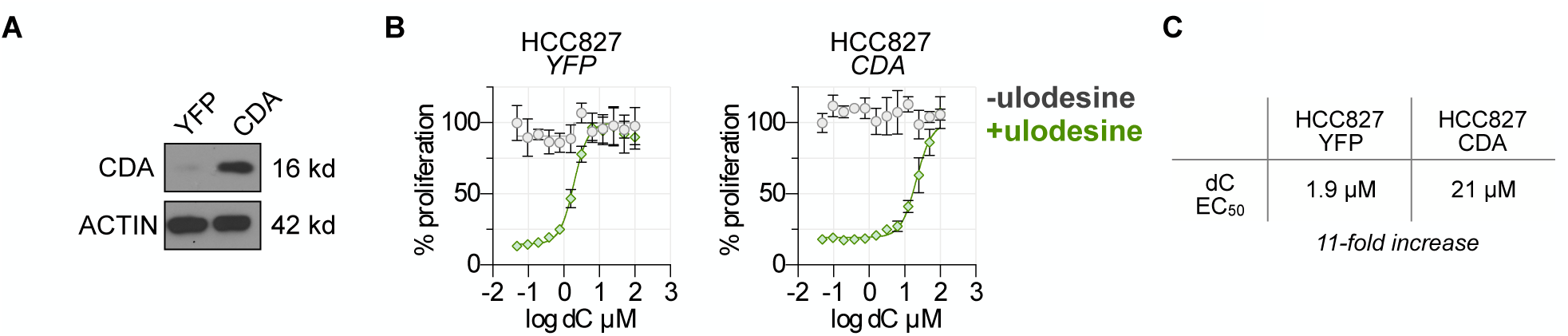
CDA overexpression mitigates competition between dC and dG and enhances ulodesine activity. (**A**) Immunoblot analysis of HCC827-YFP and -CDA over-expressing isogenic cells. (**B**) Cell Titer Glo analysis of solid cancer cell lines HCC827 isogenic cells treated with 5 µM dG ±1 µM ulodesine ±dC for 72 h (mean±SD; n=4). (**C**) EC_50_ values for dC from experiment in **C**.

## DISCUSSION

Synthetic lethality refers to the scenario where impaired function of either of two genes alone is tolerated where simultaneous loss results in cell death^32^. This concept has been explored extensively in the context of anti-cancer therapy with the goal of identifying synthetic lethal partners with cancer-associated genomic alterations. Perhaps the most well-studied example is the acute sensitivity of BRCA1-deficient cancer cells to PARP inhibitors^33^. Here we add to the library of synthetic lethal interactions in cancer and identify PNP activity as a targetable co-dependency of cells for deficient for SAMHD1.

Mutational inactivation of PNP has been linked linked to a remarkably selective T-cell deficiency phenotype in humans. We propose that the lack of SAMHD1 expression in developing thymocytes, and not dCK expression alone, underlies this selectivity. Interestingly, similar tissue selectively was observed in dCK knockout mice and has been linked to an imbalanced dTTP pool suggesting that SAMHD1 may play a role in this phenotype^30^. SAMHD1 has been shown to be expressed in mature T-cells but the reasons for its inactivation in developing hematopoietic tissues are unclear^34^. This altered expression may be linked to the high-rate of proliferation, and high dNTP demand, of these developing cells. It is likely that ALL cells are progeny of this immature T-cell population. An important corollary of our work is the finding that pharmacological dCK inhibition mitigates the phenotypes associated with PNP inactivation which suggests that dCK inhibitors could have utility in the treatment of PNP or ADA-linked SCID.

Interestingly, dGTP functions not only as a substrate by also as an allosteric activator of SAMHD1. SAMHD1 possesses two nucleotide-binding allosteric regulatory sites which effectively tune its function. Allosteric site 1 binds either GTP or dGTP and allosteric site 2 binds either dGTP, dATP, dCTP or dTTP. While the species of allosteric site-bound nucleotides influences the K_m_ of SAMHD1 for its substrate, the K_m_ for dGTP is lowest regardless of the combination of bound regulatory nucleotides^24^. Taken together, these findings suggest that a primary function of SAMHD1 is to sense and restrict any increases in dGTP pool size.

We found that PNP inhibition triggers replication stress, DNA damage and apoptosis in SAMHD1-deficient cells. However, the mechanistic link between dGTP pool expansion and induction of phenotypes remains incompletely defined. The lethal effects of dGTP pool imbalance could result dNTP insufficiency resulting from allosteric inhibition of pyrimidine nucleotide reduction by RNR^15^. Small molecule inhibitors of ATR, including M6620 and AZD6738, are currently being investigated in phase I/II trials against solid tumors and may represent rational companions for PNP inhibitors^35^. Furthermore, as SAMHD1-deficiency is requirement for PNP activity in cancer cells the development of SAMHD1 inhibitors would broaden the patient population that could be treated with forodesine or ulodesine. However, broad non-specific toxicity is a potential limitation of this approach and it remains to be determined whether a sufficient therapeutic window exists for combinations of PNP and SAMHD1 inhibitors.

Several nucleoside analog PET probes have been developed and are routinely used in the clinic in the oncology setting. The purine nucleoside analog _18_F-CFA may be particularly useful in identifying tumors most likely to accumulate dGTP following PNP inhibition as it is sensitive to alterations in the expression of both CDA and dCK_36_.

SAMHD1 has been shown to regulate multiple biological processes, including: retrovirus (HIV) restriction, cGAS/STING activation and therapeutic efficacy of nucleoside-analog chemotherapies such as cytarabine in acute myeloid leukemia^37–39^. Here, we extend these findings by demonstrating that SAMHD1-prevents lethal imbalances among natural dNTP pools and identify SAMHD1 expression as a biomarker for PNP inhibitor activity.

As PNP inhibitors have demonstrated excellent bioavailability in humans our work strongly supports reigniting their evaluation in biomarker-driven anti-cancer trials. Collectively, our results indicate that SAMHD1, CDA and dCK expression are requisite biomarkers that must be taken into consideration in any future clinical trials evaluating PNP inhibitors. Additionally, our findings challenge the dogma that disease type (i.e. T-cell leukemia/lymphoma) predicts PNP inhibitor activity and provides evidence that PNP inhibitors may have utility in the treatment of a precisely defined subset of both hematological cancers and solid tumors.

## METHODS

### Experimental Model and Subject Details

#### Cell culture

All cell cultures were between passages 3 and 20 and maintained in antibiotic free DMEM or RPMI +10% dialyzed FBS, at 37°C in 5% CO_2_. We routinely monitored for mycoplasma contamination using the PCR-based Venor Mycoplasma kit. PDAC cell lines were acquired either from a commercial vendor (ATCC, DSMZ) or from collaborators. Cell line identity was independently authenticated by PCR.

#### Drugs

Drug stocks were prepared in DMSO or H_2_O and diluted fresh in cell culture media for treatments.

### Method Details

#### Propidium iodide cell cycle analysis

Treated cells were washed with PBS and suspended in propidium iodide cell cycle staining solution (100 µg/ml propidium iodide; 20 µg/ml Ribonuclease A). 10,000 events were collected per sample. All flow cytometry data were acquired on five-laser BD LSRII, and analyzed using FlowJo software (Tree Star).

#### Viability analysis

For CellTiter-Glo analysis cells were plated at 1×10^3^ cells / well in 50 µl / well in white opaque 384-well plates and treated as described. Following incubation 50 µl of CellTiter-Glo reagent (Diluted 1:5 in dH_2_O) was added to each well, plates incubated at room temperature for 5 m and luminescence was measured using a BioTek microplate luminescence reader. Proliferation rate normalized growth inhibition was calculated using the previously described GR metric ^40^.

For crystal violet staining, cells were plated in 6-well cell culture plates at 1×10^4^ cells/well and treated as described. Following treatment cells were fixed by incubating in 4% PFA in PBS for 15 m at room temperature. Plates were subsequently washed with PBS and stained with 0.1% crystal violet in H_2_O for 15 m at room temperature.

#### Immunoblot analysis

PBS-washed cell pellets were resuspended in cold RIPA buffer supplemented with protease and phosphatase inhibitors. Protein lysates were normalized using BCA assay, diluted using RIPA and 4x laemmli loading dye, resolved on 4-12% Bis-Tris gels and electro-transferred to nitrocellulose membranes. After blocking with 5% nonfat milk in TBS + 0.1% Tween-20 (TBS-T), membranes were incubated overnight in primary antibodies diluted (per manufacturers instructions) in 5% BSA in TBS-T. Membranes were washed with TBST-T and incubated with HRP-linked secondary antibodies prepared at a 1:2500 dilution in 5% nonfat dry milk in TBS-T. HRP was activated by incubating membranes with a mixture of SuperSignal Pico and SuperSignal Femto ECL reagents (100:1 ratio). Exposure of autoradiography film was used for detection.

#### Retroviral transduction and stable cell line generation

The pMSCV-hCDA-IRES-EYFP plasmid was described previously^41^. Amphotropic retroviruses were generated by transient co-transfection of the MSCV retroviral plasmid and pCL-10A1 packaging plasmid into Phoenix-Ampho packaging cells.

#### Generation of SAMHD1 knockout cells using CRISPR/Cas9

A SAMHD1-targeting gRNA(TCTCGGGCTGTCATCGCAAC) was cloned into the lentiCRISPRv2 backbone and used for transfection of SUIT2 cells using Lipofectamine3000. Following transfection, SUIT2 cells were selected with 5 µg/mL puromycin for 72 h and singly cloned in 96-well plates. Knockout clones were validated using immunoblot analysis.

### Key Resource Table

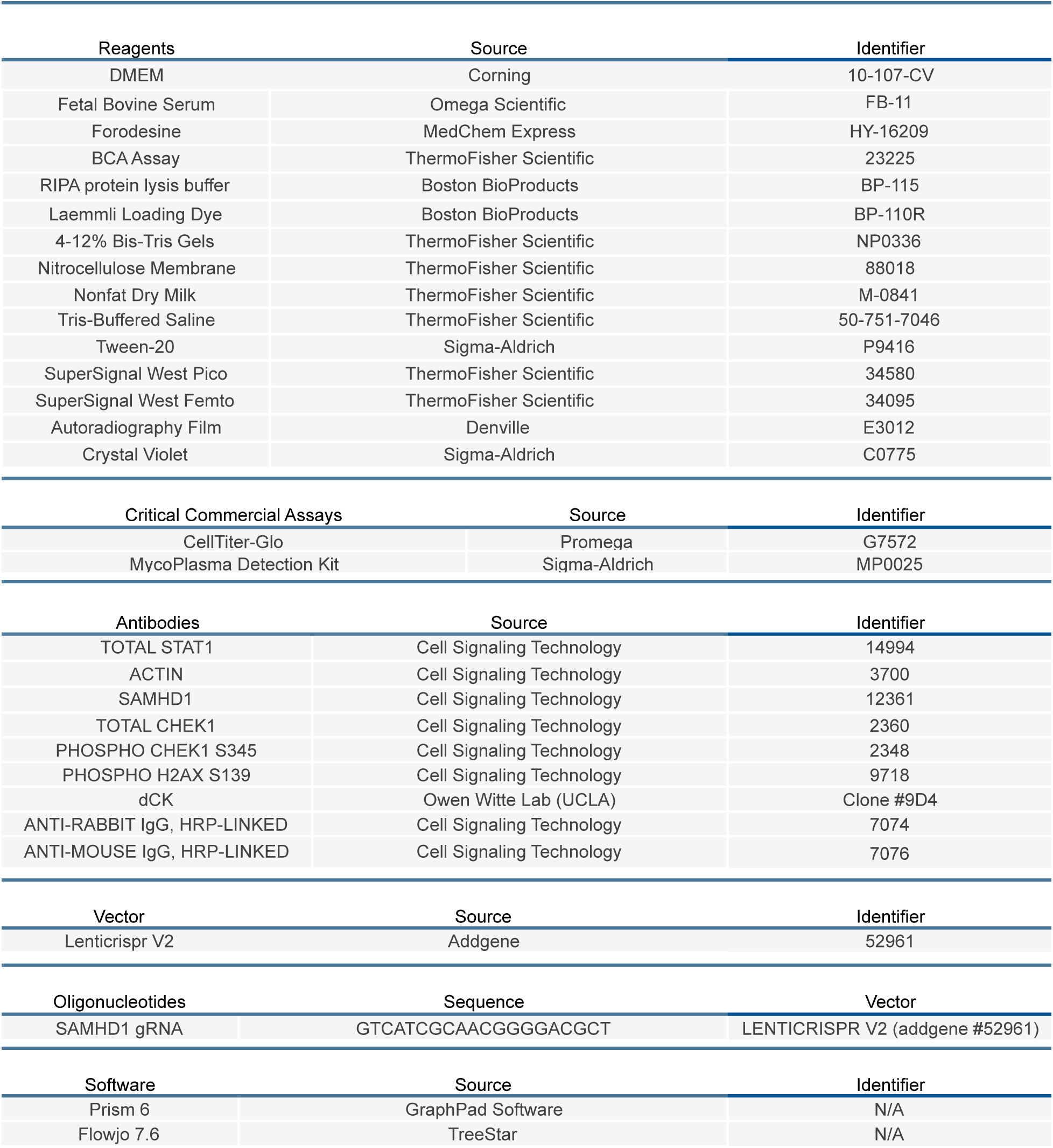

